# Cell-autonomous regulation of epithelial cell quiescence by calcium channel Trpv6

**DOI:** 10.1101/631861

**Authors:** Yi Xin, Allison Malick, Meiqin Hu, Chengdong Liu, Heya Batah, Haoxing Xu, Cunming Duan

**Affiliations:** Department of Molecular, Cellular and Developmental Biology, University of Michigan, Ann Arbor, MI 48109

**Author notes:** Corresponding to: Dr. Cunming Duan, Department of Molecular, Cellular, and Developmental Biology, University of Michigan, Ann Arbor, MI 48109.

**Keywords:** TRPV6, Ca^2+^ signaling, IGF1 receptor, Akt, mTor, PP2A, zebrafish, colon cancer cell

## Abstract

Epithelial homeostasis and regeneration require a pool of quiescent cells. How the quiescent cells are established and maintained is poorly understood. Here we report that Trpv6, a cation channel responsible for epithelial Ca^2+^ absorption, functions as a key regulator of cellular quiescence. Genetic deletion and pharmacological blockade of Trpv6 promoted zebrafish epithelial cells to exit from quiescence and re-enter the cell cycle. Reintroducing Trpv6, but not its channel dead mutant, restored the quiescent state. Ca^2+^ imaging showed that Trpv6 is constitutively open in vivo. Mechanistically, Trpv6-mediated Ca^2+^ influx maintained the quiescent state by suppressing insulin-like growth factor (IGF)-mediated Akt and Tor signaling. In zebrafish epithelia and human colon cancer cells, Trpv6/TRPV6 elevated intracellular Ca^2+^ levels and activated PP2A, which down-regulated IGF signaling and promoted the quiescent state. Our findings suggest that Trpv6 mediates constitutive Ca^2+^ influx into epithelial cells to continuously suppress growth factor signaling and maintain the quiescent state.

## Introduction

Quiescence is a non-proliferative cellular state found in many cell types in the body. While non-proliferative, these cells retain the ability to re-enter the cell cycle in response to appropriate cell-intrinsic and extrinsic signals (Matson & Cook, 2017; Sun & Buttitta, 2015; Yao, 2014). Quiescence protects long-lived cells, such as adult stem cells against the accumulation of genomic aberrations and stress. Maintaining a pool of quiescent cells is critical for tissue repair, wound healing, and regeneration (Cheung & Rando, 2013). This is particularly important for epithelia which are rapidly and continuously renewed throughout life. The intestinal epithelial cells, for example, are renewed every 4 to 5 days (van der Flier & Clevers, 2009). By synchronizing cultured mammalian cells in G0 via serum starvation followed with serum re-stimulation, Yao et al. (2008) showed that the Rb proteins (pRb, p107, and p130) and their interactions with E2F proteins are critical in regulating the proliferation-quiescence decision. Acting downstream, a bifurcation mechanism controlled by CDK2 activity and p21 regulating the proliferation-quiescence decision has also been demonstrated in cultured mammalian cells (Spencer et al., 2013). While important insights have been learnt from in vitro studies, how the quiescent cell pools are established during development and maintained in vivo is not well understood. The exceptionally high turnover rate implies that cell type-specific mechanism(s) must exit.

The transient receptor potential cation channel subfamily V member 6 (TRPV6) is expressed in mammalian intestinal epithelial cells (Hoenderop et al., 2005). TRPV6 is a conserved calcium channel that constitutes the first and rate-limiting step in the transcellular Ca^2+^ transport pathway (Hoenderop et al., 2005; Peng et al., 1999; Peng et al., 2000; Wissenbach et al., 2001). In zebrafish, *trpv6* is specifically expressed in a population of epithelial cells known as ionocytes or NaR cells (Dai et al., 2014; Pan et al., 2005). NaR cells take up Ca^2+^ from the surrounding habitats into the body to maintain body Ca^2+^ homeostasis (Liao et al., 2009). NaR cells are polarized cells that functionally and molecularly similar to human intestinal epithelial cells. While located in the gill filaments and the intestine in the adult stages, these cells are distributed in the yolk sac skin during the embryonic and larval stages, making these easily accessible for experimental observation and perturbations (Dai et al., 2014; Pan et al., 2005). When zebrafish are grown in homeostatic normal [Ca^2+^] conditions, NaR cells are maintained in a quiescent state and the Akt-Tor activity is regulated at low levels. Low [Ca^2+^] stress increases Akt-Tor activity in these cells and promotes their re-entry into the cell cycle (Dai et al., 2014; Liu et al., 2017). This is similar to the proposed role of mTOR signaling in adult stem cells (Kim & Guan, 2019; Meng et al., 2018), suggesting an evolutionarily conserved mechanism(s) at work. More recent studies suggest that insulin-like growth factor binding protein 5a (Igfbp5a), a secreted protein that binds IGF with high-affinity, plays a critical role in activating Akt-Tor signaling in these cells via the IGF1 receptor under calcium deficient states (Liu et al., 2018). The mechanism controlling the quiescent state under normal [Ca^2+^] condition is currently unknown. In a previous study, we found that zebrafish *mus* mutant larvae, a loss-of-function Trpv6 mutant fish line obtained from an ENU mutagenesis screen (Vanoevelen et al., 2011), had many proliferating NaR cells and elevated Akt-Tor signaling, suggesting Trpv6 may play a negative role in regulating NaR cell proliferation (Dai et al., 2014). How does Trpv6 act to inhibit Akt-Tor signaling and whether it involves in quiescence regulation are unknown. Because TRPV6/Trpv6 is the primary Ca^2+^ channel responsible for epithelial Ca^2+^ uptake and since Ca^2+^ is a major second messenger involved in cell proliferation and differentiation in many cell types (Clapham, 1995; Hoenderop et al., 2005), we hypothesized that Trpv6 regulates the quiescent state by conducting Ca^2+^ influx into epithelial cells and suppressing IGF1 receptor mediated Akt-Tor signaling. The objective of this study was to test this hypothesis and to elucidate the underlying mechanisms of Trpv6 action.

## Results

### Trpv6 is crucial for epithelial Ca^2+^ uptake in zebrafish

Three *trpv6* mutant fish lines were generated using CRISPR/Cas9 (Fig. 1A). All three Trpv6 mutant proteins lack the 6 transmembrane domains and the critical ion pore region and are predicted to be null mutations (Fig. 1B). The *trpv6Δ7* and *trpv6Δ8* lines were made in the *Tg(igfbp5a:GFP)* fish background. *Tg(igfbp5a:GFP)* is a transgenic fish line expressing EGFP in the *trpv6*-expressing NaR cells (C. Liu et al., 2017), allowing real-time analysis of NaR cell proliferation. The *trpv*6*Δ8-2* line was in a non-transgenic fish background and used in Ca^2+^ imaging analysis described later. The gross morphology and body size of the mutant fish were similar to their siblings (Fig. S1). All mutant fish died within 2 weeks (Fig. 1C and 1D). Alizarin red staining indicated a marked reduction in the calcified bone mass in the *trpv6*^*-/-*^ mutant fish (Fig. 1E), indicating body calcium deficiency. Fura-2 Ca^2+^ imaging experiments using HEK293 cells transfected with zebrafish Trpv6 were performed. The Trpv6-mediated [Ca^2+^]_i_ change was similar to that of human TRPV6 (Fig. 1F). D542 in mammalian TRPV6 occupies a critical position in the ion pore region and mutation of this residue abolishes its Ca^2+^ permeability (McGoldrick et al., 2018; Sakipov, Sobolevsky, & Kurnikova, 2018). This residue is conserved in zebrafish Trpv6 at position 539 (Fig. S2). We generated and tested Trpv6D539A mutant. The [Ca^2+^]_i_ levels in Trpv6D539A mutant transfected cells were low and did not respond to changes in extracellular [Ca^2+^] (Fig. 1F). The maximal Ca^2+^ influx rate was reduced to a negligible level in Trpv6D539A transfected cells (Fig. 1G). Whole-cell patch clamp experiments confirmed that the Trpv6 mediated Ca^2+^ current and this activity was abolished in the Trpv6D539A mutant (Fig. 1H). These findings support the notion that Trpv6 play an indispensable role in epithelial Ca^2+^ uptake and maintaining body Ca^2+^ balance and provided critical reagents for subsequent experiments.

**Fig. 1.**
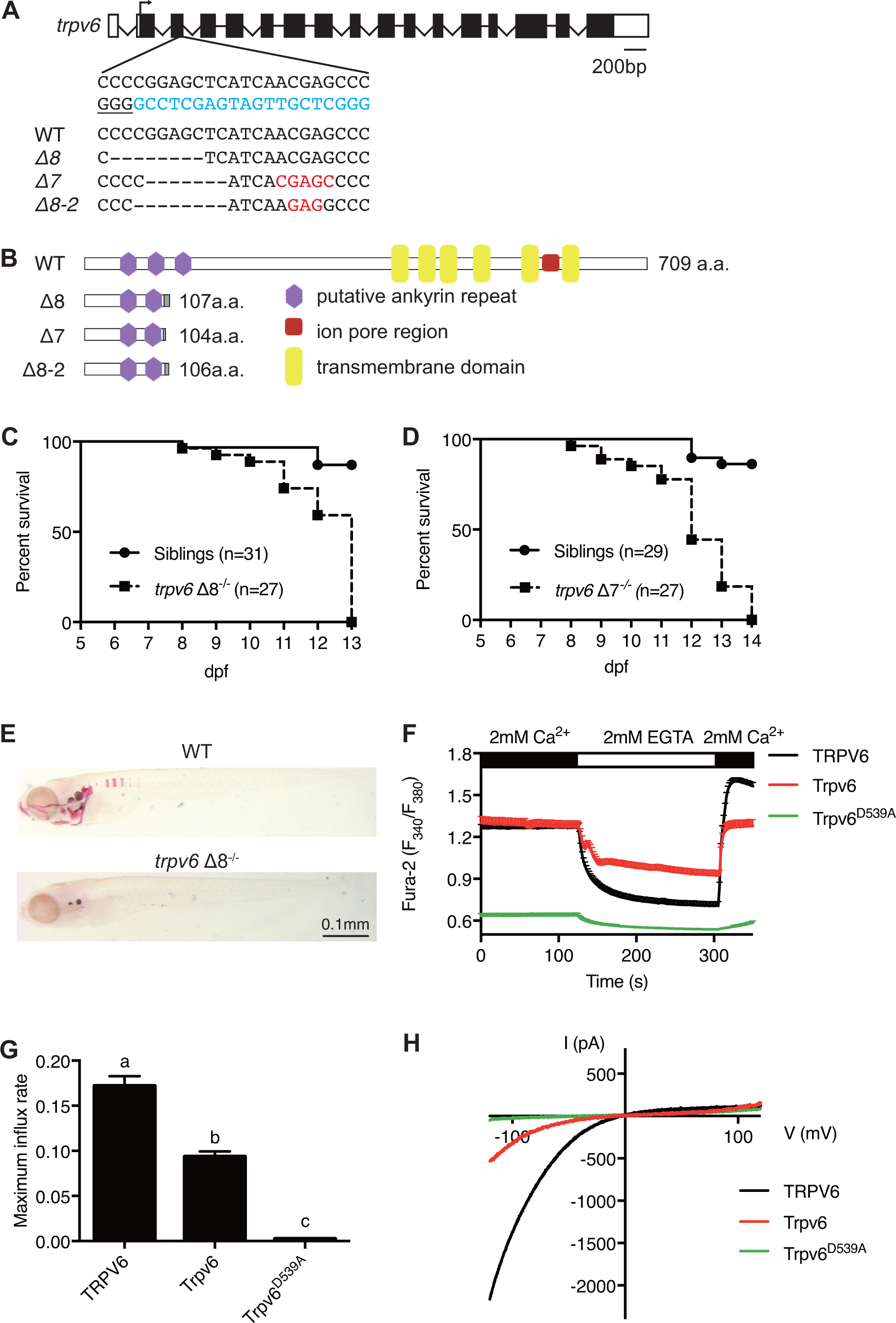
Genetic deletion of the conserved epithelial calcium channel Trpv6 results in calcium deficiency and premature death. (**A**) Schematic diagram showing *trpv6* gene (WT) and various mutant sequence. Filled boxes indicate *trpv6* ORF and open boxes indicate UTRs. Introns are shown as lines. The gRNA targeting site is indicated by blue color and PAM motif is underlined. Deleted and inserted nucleotides are indicated by dash lines and red letters, respectively. (**B**) Schematic diagram of Trpv6 (WT) and its mutants. Putative function domains are indicated. The grey box indicates altered sequence caused by frame shifts. (**C-D**) Survival curves of *trpv6Δ8*^*-/-*^; *Tg(igfbp5a:GFP)* (**C**) and *trpv6Δ7*^*-/-*^; *Tg (igfbp5a:GFP)* fish (**D**) and siblings. The numbers of total fish are indicated. (**E**) Representative images of Alizarin red stained wild-type and *trpv6Δ8*^*-/-*^; *Tg(igfbp5a:GFP)* fish at 7 day post fertilization (dpf). (**F**) Fura-2 Ca^2+^ imaging analysis of HEK293 cells transfected with the indicated plasmids. n > 50 cells from 3 independent experiments. (**G**) The maximal influx rate. n = 3 independent experiments. (**H**) Currents evoked by a RAMP voltage from −120mV to +120 mV in HEK293 cells transfected with the indicated plasmids. In this and all subsequent figures, unless specified otherwise data shown are Mean ± SEM. Different letters indicate significant difference at *P* < 0.05, one-way ANOVA followed by Tukey’s multiple comparison test.

### Trpv6 regulates the quiescence-proliferation decision in epithelial cells

To determine the possible role of Trpv6 in NaR cells, double-blind tests were performed (Fig. 2A). In agreement with previous studies (Dai et al., 2014; Liu et al., 2017), NaR cells in the wild-type and heterozygous siblings were distributed in the yolk sac region as single cells in a salt-and-pepper pattern (Fig. 2B). NaR cells in the *trpv6Δ8* mutant larvae were often observed in clusters of newly divided cells (Fig. 2B). These proliferating NaR cells had enlarged apical opening (Fig. S3). The average NaR cell proliferation rate was elevated in the mutant fish in all stages examined (Fig. 2C). At 5 dpf, the *trpv6Δ8* mutant fish had 3-time more NaR cells (Fig. 2C). Essentially same data were obtained with the *trpv6Δ7* fish (Fig. 2D). GdCl_3_, a Trpv6 inhibitor, was used to further test the role of Trpv6. GdCl_3_ treatment increased NaR cell proliferation in the wild-type and heterozygous fish, while it did not further increase NaR cell proliferation in the mutant fish (Fig. 2E). Ruthenium red, another Trpv6 inhibitor, had similar effects (Fig. S4). Next, Trpv6 and Trpv6D539A were randomly expressed in NaR cells in *trpv6Δ8*^*-/*−^; *Tg(igfbp5a:GFP)* fish using a Tol2 transposon BAC-mediated genetic mosaic assay (Chengdong Liu et al., 2018). Reintroduction of Trpv6 reversed the quiescence to proliferation transition (Fig. 2F) and reduced the apical opening site to the control levels (Fig. S3). Trpv6D539A, however, had no such effect (Fig. 2F). These data showed that Trpv6 functions as a major barrier in the quiescence to proliferation transition and this action requires its Ca^2+^ permeability.

**Fig. 2.**
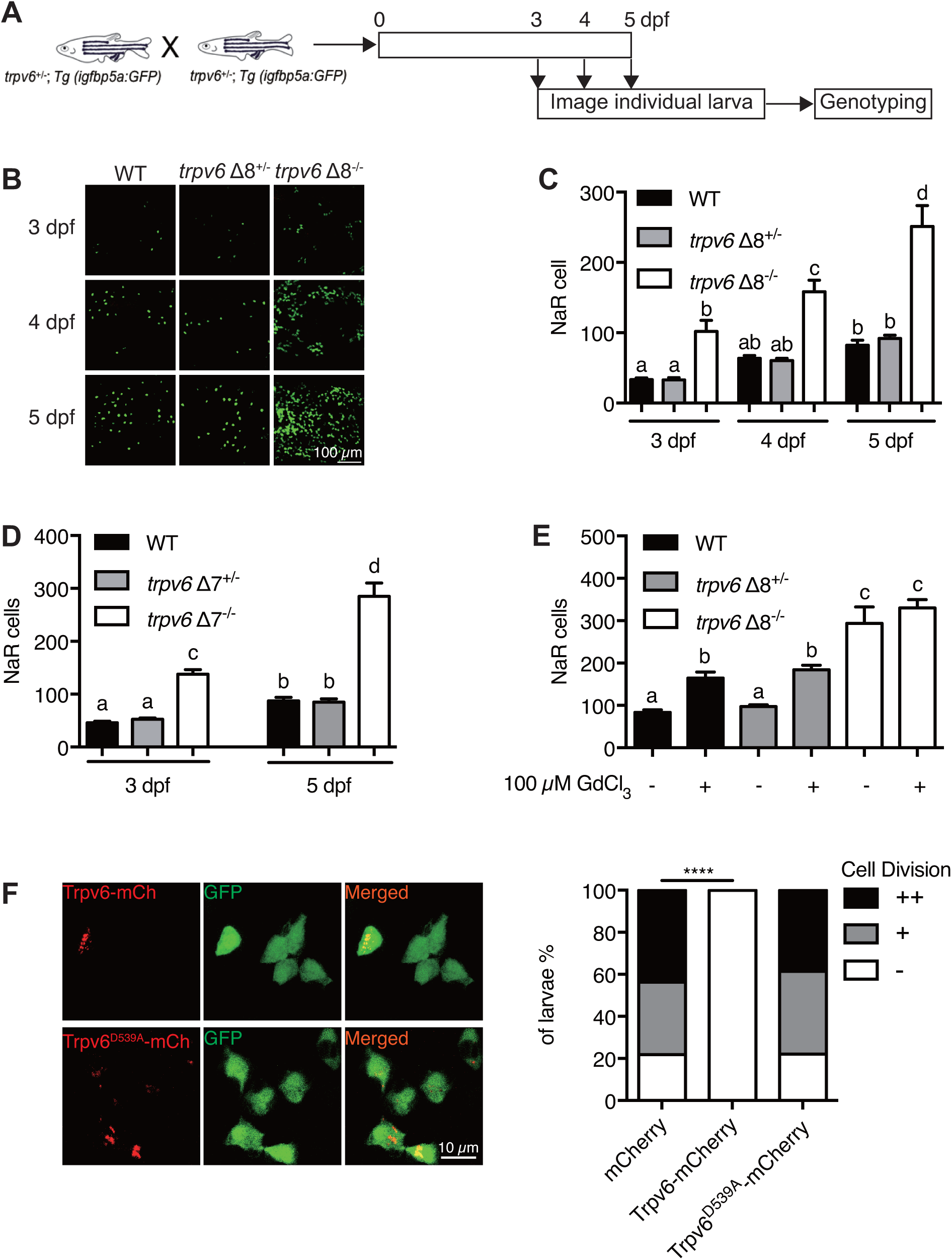
Trpv6 regulates epithelial cell quiescence-proliferation decision. (**A**) Diagram of the experimental design. (**B**) Representative images. In this and all subsequent larval images, lateral views of the yolk-sac region are shown with dorsal up and anterior to the left. (**C-D**) Mean NaR cell number/fish of the indicated genotypes. n = 6-9. (**E**) Progenies of *trpv6 Δ8*^*+/-*^; *Tg (igfbp5a:GFP)* intercross were raised to 3 dpf and treated with 100 µM GdCl_3_ from 3 to 5 dpf. NaR cells in each fish were quantified following individual genotyping. n = 13-22. (**F**) Progenies of *trpv6Δ8*^*+/-*^*;Tg(igfbp5a:GFP)* intercross were injected with the indicated BAC-mCherry DNA at one-cell stage. At 5 dpf, the Trpv6-expressing NaR cells in each fish were scored following a published scoring system (Chengdong Liu et al., 2018). Representative images are shown in the left and quantified results in the right panel. ****, *P* <0.0001 by Chi-Square test, fish n = 12-38.

### Trpv6 controls the quiescence-proliferation decision via regulating growth factor signaling

Previous studies in wild-type fish suggested that NaR cells re-enter the cell cycle in response to low [Ca^2+^] treatment (Dai et al., 2014; Liu et al., 2017). To determine whether this effect is related to Trpv6, 3 dpf *trpv6Δ7*^*-/-*^*;Tg(igfbp5a:GFP)* larvae and siblings were subjected to low [Ca^2+^] challenge test. Low [Ca^2+^] treatment for 2 days resulted in a 3-fold increase in proliferating NaR cells in the wild-type and heterozygous fish (Fig. 3A). This value was comparable to that of *trpv6Δ7*^*-/-*^ larvae kept in normal [Ca^2+^] condition (Fig. 3A). Low [Ca^2+^] treatment did not further increase NaR cell number in the mutant larvae (Fig. 3A). Immunostaining results showed significant increases in the number of phosphorylated Akt-positive NaR cells in *trpv6Δ7*^*-/-*^ *and trpv6Δ8*^*-/-*^ larvae kept in the normal [Ca^2+^] embryo solution (Fig. 3B-3D). In comparison, the levels of phospho-Akt in the siblings were minimal. Tor signaling activity was also significantly elevated in the mutant larvae (Fig. 3E-3G). Blocking Trpv6 channel activity using GdCl_3_ and Ruthenium red increased phospho-Akt levels and NaR cell proliferation in the wild-type fish (Fig. S4). If Trvp6 regulates the quiescence-proliferation decision via suppressing the IGF1 receptor-mediated Akt signaling, then reintroduction of Trpv6 should inhibit Akt signaling and blockade of IGF signaling should reverse the quiescent to proliferation transition. Indeed, double label staining showed that re-expression of Trpv6 inhibited Akt phosphorylation in NaR cells (Fig. 3H). Treatment of the mutant fish with BMS-754807, an IGF1 receptor inhibitor, abolished the quiescence to proliferation transition (Fig. 3I-3J).

**Fig. 3.**
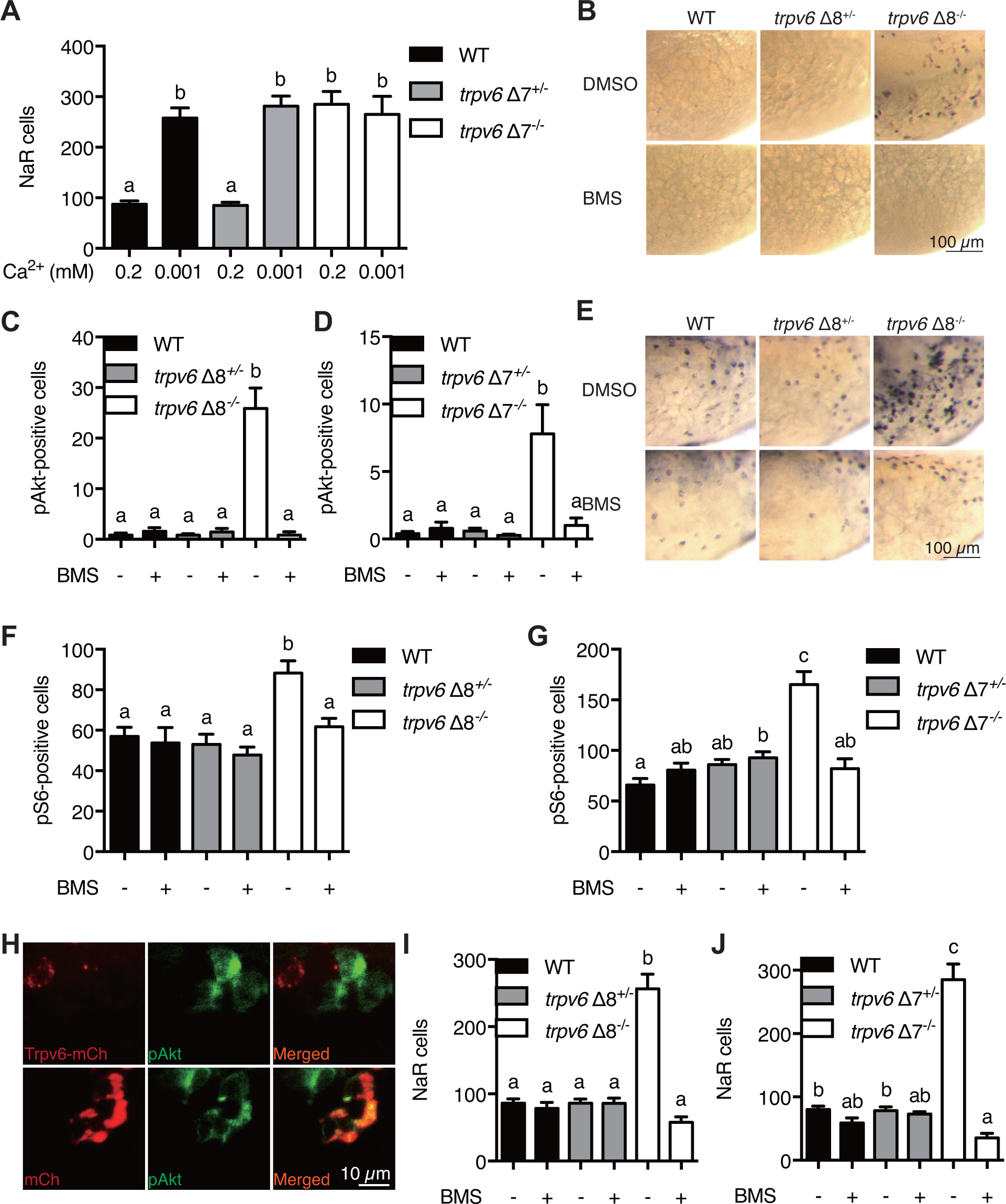
Trpv6 prevents the quiescence to proliferation transition via regulating IGF1 receptor-mediated Akt and Tor signaling. **(A)** Progenies of *trpv6*Δ7^*+/-*^; *Tg(igfbp5a:GFP)* intercrosses were grown in embryo solutions with the indicated Ca^2+^ concentration from 3 dpf to 5 dpf. NaR cells in each fish were quantified followed by individual genotyping. n = 5-17 fish. (**B-G**) Embryos of the indicated genotypes were raised to 3 dpf and treated with 0.3 µM BMS-754807 or DMSO. At 4 dpf, the treated fish were subjected to immunostaining using an anti-phospho-Akt antibody (**B-D**) or an anti-phospho-S6 antibody (**E-G**). Representative images are shown in (**B** and **E**). Quantified data are shown in (**C, D, F** and **G**). n = 5-41. (**H**) Progenies of a *trpv6Δ8*^*+/-*^; *Tg (igfbp5a:GFP)* intercross were injected with the indicated BAC-mCherry DNA at one-cell stage. At 4 dpf, the larvae were subjected to phospho-Akt and mCherry double staining. (**I-J**) Progenies of *trpv6*Δ8^*+/-*^; *Tg(igfbp5a:GFP)* (**I**) or *trpv6*Δ7^*+/-*^; *Tg(igfbp5a:GFP)* (**J**) intercrosses were raised to 3 dpf and treated with BMS-754807 or DMSO from 3 to 5 dpf. NaR cells in each fish were quantified followed by individual genotyping, n = 6-22.

### Trpv6 constitutively conducts Ca^2+^ into epithelial cells and regulates the [Ca^2+^]_i_ levels in vivo

Although it is well accepted that TRPV6 is the primary epithelial Ca^2+^ channel, all reported electrophysiology studies to date were done in cultured mammalian cells over-expressing TRPV6. TRPV6-mediated Ca^2+^ influx or current has not been recorded in vivo or even in primary epithelial cells (Fecher-Trost, Wissenbach, & Weissgerber, 2017). To investigate Trvp6-mediated Ca^2+^ influx in vivo, we generated the *Tg(igfbp5a:GCaMP7a)* fish, a stable reporter fish line expressing GCaMP7a in NaR cells (Fig. S5A). After validating the effectiveness of GCaMP7a in reporting intracellular Ca^2+^ levels ([Ca^2+^]_i_) (Fig. S5B and S5C), *trpv*6*Δ8-2*^+/−^; *Tg(igfbp5a:GCaMP7a)*^*+/-*^ fish were crossed with *trpv*6*Δ8-2*^+/−^ fish and their offspring were screened at 3 dpf and then genotyped individually. While GCaMP7a-positive cells were observed in ∼50% of the siblings as expected, none of the *trpv*6*Δ8-2*^*-/-*^ mutant larvae had any GCaMP7a-positive cells (Fig. 4A and 4B). Addition of the Ca^2+^ ionophore ionomycin restored GCaMP7a signal in the mutant fish to a level comparable to their siblings (Fig. 4B and 4C), thus ruling out the possibility that GCaMP7a expression is altered in the mutant fish. Next, Trpv6 was randomly expressed in NaR cells in the *trpv6 Δ8-2*^*-/*−^; *Tg(igfbp5a:GCaMP7a)* fish using the genetic mosaic assay. Reinduction of Trpv6 significantly increased GCaMP7a signal levels (Fig. 4D). Ionomycin treatment did not result in further increase (Fig. 4D). These genetic data argue strongly that Trpv6 is not only critical in conducting Ca^2+^ into NaR cells, but also in maintaining the high [Ca^2+^]_i_ levels in these cells. We next used Trpv6 inhibitors to block Trpv6 activity in *Tg(igfbp5a:GCaMP7a)* fish. Within 8 min after the GdCl_3_ treatment, the [Ca^2+^]_i_ levels became significantly lower and the reduction became more pronounced in 12 and 16 min (Fig. 4E and 4G; Video S1). When GdCl_3_ was washed out, the [Ca^2+^]_i_ levels gradually increased and returned to normal levels after 12 min (Fig. 4F and 4G; Video S2). Similar results were obtained with Ruthenium red (Fig. S5D-S5E). Addition of the IGF1 receptor inhibitor BMS-754807 or PP2A inhibitors Okadaic acid or Calyculin did not change the [Ca^2+^]_i_ levels in NaR cells (Fig. S6A-S6F). Therefore, Trpv6 constitutively conducts Ca^2+^ into epithelial cells and continuously maintains high [Ca^2+^]_i_ levels in vivo.

**Fig. 4.**
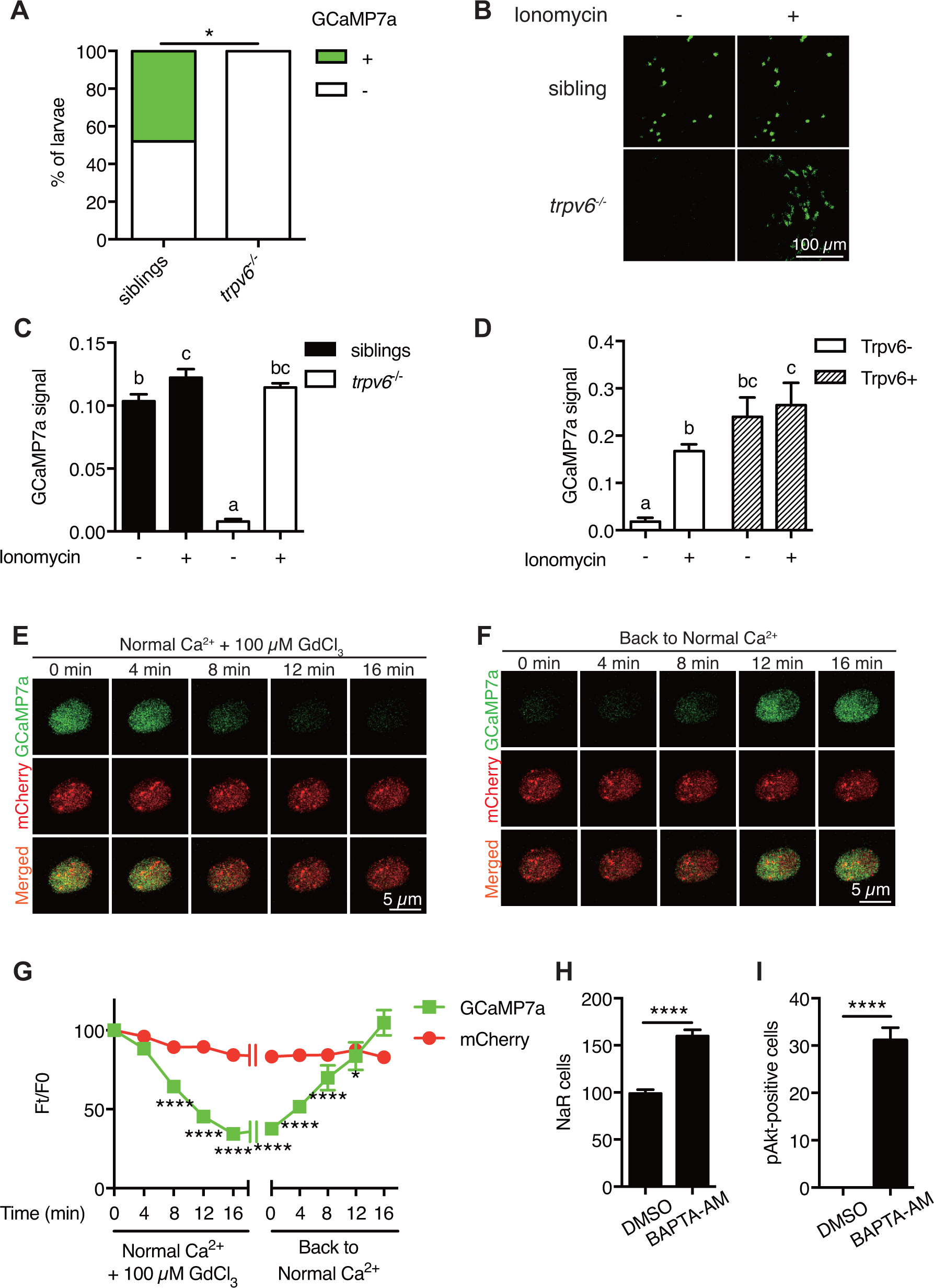
Trpv6 is constitutively open and mediates Ca^2+^ influx and maintain high [Ca^2+^]_i_ in epithelial cells in vivo. (**A**) *trpv6Δ8-2*^*+/-*^*;Tg (igfbp5a:GCaMP7a)*^*+/-*^ was crossed with *trpv6Δ8-2*^*+/-.*^ The progenies were imaged at 3 dpf followed by individual genotyping. Percentage of GCaMP7a-positive fish is shown. *, *P* < 0.05 by Chi-Square test, n = 22. (**B-C**) Fish described in (**A**) were imaged before and after the addition of 7.5 µM Ionomycin + 10 mM CaCl_2_. Representative images are shown in (**B**) and the quantified results are shown in (**C**). n = 5-7. (**D**) Progenies from a *trpv6Δ8-2*^*+/-*^*;Tg (igfbp5a:GCaMP7a)*^*+/-*^ and *trpv6Δ8-2*^*+/-*^ intercross were injected with *BAC (igfbp5a:Trpv6-mCherry)* DNA at 1-cell stage. They were raised to 3 dpf. GCaMP7a signal intensity in Trpv6-mCherry-expressing cells and non-expressing NaR cells were quantified before and after the addition of 7.5 µM Ionomycin+10 mM CaCl_2_. n = 4. (**E-G**) Time-lapse images of 3 dpf *Tg (igfbp5a:GCaMP7a)* larvae after the addition of 100 µM GdCl_3_ (**E**) or following drug removal (**F**). Changes in GCaMP7a and mCherry signal intensity ratio were quantified and show in (**G**). n = 5. * and **** indicate *P* < 0.05 and < 0.0001 by Two-way ANOVA followed by Dunnett’s multiple comparisons test. (**H**) Wild-type larvae were treated with BAPTA-AM (100 µM) from 3 dpf to 5 dpf. NaR cells were labeled by *in situ* hybridization using a *trpv6* riboprobe and quantified. (**I**) Larvae described in (**H**) were stained for phosphorylated Akt after 24 hour treatment. Mean ± SEM. ****, *P*<0.0001, unpaired t-test. n = 15-19.

### Trpv6 inhibits IGF signaling and epithelial cell proliferation by regulating [Ca^2+^]_i_ and PP2A is a downstream effector

The observation that [Ca^2+^]_i_ in NaR cells are continuously maintained at high levels was surprising and intriguing. To determine whether the observed high [Ca^2+^]_i_ levels have any functional significance, *Tg(igfbp5a:GFP*) larvae were treated with the intracellular Ca^2+^ chelator BAPTA-AM. BAPTA-AM treatment resulted in a significant increase in NaR cell number (Fig. 4H) and in phospho-Akt signaling levels (Fig. 4I). These data indicate that the high [Ca^2+^]_i_ levels are critical in regulating the quiescent state. To identify the downstream effector(s) of [Ca^2+^]_i_, a collection of small molecule inhibitors with known protein targets were screened using *Tg(igfbp5a:GFP*) larvae. Okadaic acid and Calyculin, two inhibitors of the conserved protein phosphatase 2A (PP2A), were among the strongest hits. Treatment of *Tg(igfbp5a:GFP*) larvae with either drug significantly increased NaR cell proliferation rate (Fig. 5A and 5B). This effect is specific because the drug treatment had no such effect in PP2A deficient zebrafish (Fig. S7C). Importantly, the Okadaic acid and Calyculin treatment-induced quiescence-proliferation transition was abolished by the IGF1 receptor inhibitor BMS-754807 (Fig. 5A and 5B). Okadaic acid or Calyculin treatment also resulted in significant increases in the phosphorylated-Akt levels in an IGF1 receptor-dependent manner (Fig. 5C). Okadaic acid or Calyculin treatment did not change the [Ca^2+^]_i_ levels (Fig. S6C-S6F), indicating that PP2A acts downstream of the [Ca^2+^]_i_.

**Fig. 5.**
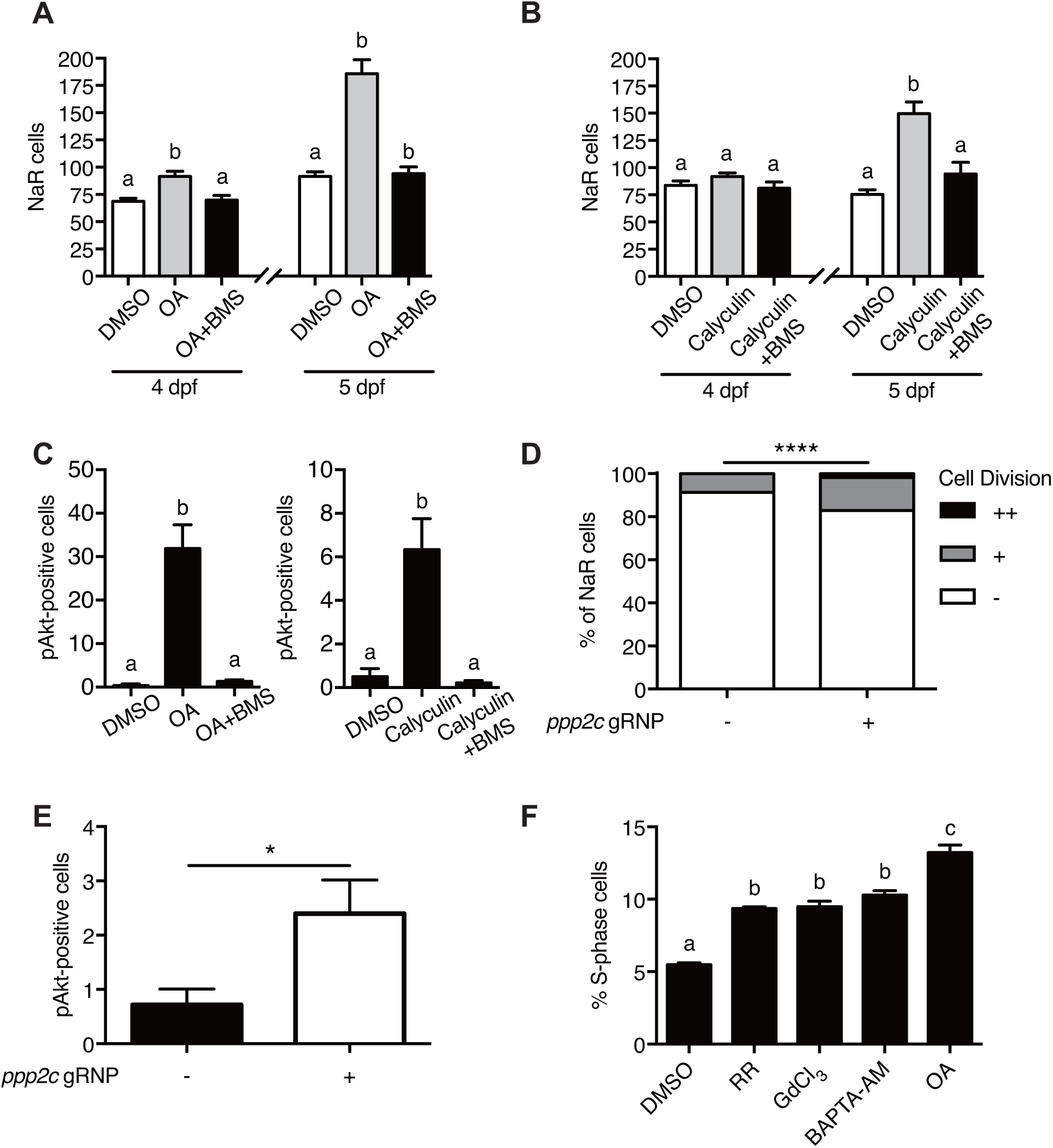
PP2A is a downstream effector of Trpv6. (**A-B**) *Tg(igfbp5a:GFP)* embryos were treated with 1 µM Okadaic acid (OA) or 0.1 µM Calyculin A in the presence or absence of 0.3 µM BMS-754807 from 3 dpf. NaR cells were quantified at 4 and 5 dpf. Data shown are n = 10-38. (**C**) Wild-type larvae were treated with 1 µM Okadaic acid or 0.1 µM Calyculin in the presence or absence of 0.3 µM BMS-754807 from 3 dpf to 4 dpf. They were analyzed by immunostaining for phospho-Akt. n = 9-16. (**D**) *Tg(igfbp5a:GFP)* embryos were injected with gRNAs targeting three *ppp2c* genes and Cas9 protein at one-cell stage. They were raised to 5 dpf. NaR cell division was quantified following a published scoring system (Chengdong Liu et al., 2018). n = 24-28. ****, *P*< 0.0001 by Chi-Square test. (**E**) The embryos treated as in (**D**) were raised to 4 dpf and analyzed by immunostaining for phospho-Akt signal. n = 20-21. *, *P*< 0.05, unpaired t-test. (**F**) Human LoVo colon cancer cells were treated with Ruthenium Red (RR, 100 µM), GdCl_3_ (100 µM), BAPTA-AM (100 µM), or Okadaic acid (OA, 20 nM) for 48 hours and analyzed by flow cytometry analysis after propidium iodide staining. n = 3.

PP2A are a family of conserved protein phosphatases that dephosphorylate Akt and many other proteins (Perrotti & Neviani, 2013; Seshacharyulu, Pandey, Datta, & Batra, 2013). PP2A holoenzymes are heterotrimers. The core enzyme is made by a catalytic C subunit (Cα and Cβ isoform), a scaffold A subunit (Aα and Aβ), and many regulatory B subunits (Virshup & Shenolikar, 2009). The combination of these subunits results in a very large number of different holoenzyme complexes. Our database search suggests that the zebrafish genome contains 3 C subunit genes (*ppp2*ca, cb, and cc). We used CRISPR/Cas9 to transiently knockdown the *ppp2c* genes because stable knockout is likely embryonic lethal. The effectiveness of the targeting guide RNAs was validated (Fig. S7A and S7B). Transient knockdown of *ppp2cs* resulted in significant increases in the number of proliferating NaR cells (Fig. 5D) and in phospho-Akt levels (Fig. 5E), suggesting that PP2A mediates the action of Trpv6-mediated Ca^2+^ influx in zebrafish epithelia. Likewise, treatment of human colon carcinoma cells with Ruthenium red, GdCl_3_, BAPTA-AM, and Okadaic acid all significantly increased cell proliferation (Fig. 5F), suggesting this regulatory axis acts in a similar manner in human colon epithelial cells.

## Discussion

In this study, we uncover a previously unrecognized role of Trpv6 and delineates a Trpv6-mediated and evolutionarily conserved Ca^2+^ signaling pathway controlling the quiescent state in epithelial cells. We showed that genetic deletion of Trpv6 not only impaired Ca^2+^ uptake and reduced body Ca^2+^ content, but also promoted epithelial cells to exit quiescence and proliferate. Likewise, pharmacological inhibition of Trpv6 increased epithelial cell quiescence-proliferation transition. While low [Ca^2+^] treatment increased epithelial cell proliferation in the siblings, it had no such effect in *trpv6*^−/−^ larvae, supporting the notion that Trpv6 functions as a major regulator of the quiescent state. Our genetic mosaic analysis results indicated that the quiescent state is regulated by Trpv6 in a cell autonomous manner. Reintroduction of Trpv6 in the *trpv6*^−/−^ mutant fish was sufficient to restore the [Ca^2+^] levels, suppress Akt-Tor signaling, and reverse the cells back to the quiescent state. This action of Trpv6 requires its Ca^2+^ conductance activity.

The impaired Ca^2+^ uptake, reduced body Ca^2+^ content, and premature death observed in *trpv6Δ7* and *trpv6Δ8* mutant fish are in good agreement with a previous study by Vanoevelen et al. (2011) using the *mus* mutant fish, but differ considerable from mouse studies. Bianco et al. reported that *Trpv6* knockout mice were viable but had reduced intestinal Ca^2+^ uptake, increased urinary Ca^2+^ excretion, decreased bone mineral density, and decreased growth and fertility (Bianco et al., 2007). Another *Trpv6*^−/−^ mutant mouse line reported by Chen et al., however, had normal blood Ca^2+^ concentration and normal bone formation, but with an increased number of osteoclasts (Chen et al., 2014). A third *Trpv6*^*-/-*^ mutant mouse model showed reduced fertility in male only (Weissgerber et al., 2012). The reason(s) of these discrepancies among these mouse studies is not fully understood, but factors such as dietary Ca^2+^ contents may have been critical (Van der Eerden et al., 2012). This may also contribute to the premature phenotypes found in the zebrafish mutants. Zebrafish embryos continuously lose Ca^2+^ and other ions into the surrounding hypo-osmotic environments and must constantly take up Ca^2+^ from the habitat to survive (Chengdong Liu et al., 2018). Another factor to take into consideration is genetic redundancy. In mammals, there is another closely related TRPV sub-family member, TRPV5. TRPV5 plays similar roles in the transcellular Ca^2+^ transport pathway, although it is mainly expressed in the kidney (Hoenderop et al., 2005). It has been shown that TRPV5 expression in the intestine is elevated in *Trpv6* mutant mice (Woudenberg-Vrenken et al., 2012). In comparison, zebrafish genome has a single trpv6 gene and lacks this genetic redundancy (Vanoevelen et al., 2011).

An important finding made in this study is that TRPV6 mediates constitutive Ca^2+^ influx into epithelial cells in vivo. Early Fura-2 Ca^2+^ imaging studies in cultured mammalian cells transfected with TRPV6 indicated that this channel may be constitutively open (Vennekens et al., 2000). This notion, however, was challenged because patch-clamp recordings using mammalian cells transfected with TRPV6 indicated no spontaneous channel activity (Bodding & Flockerzi, 2004). It was suggested that the elevated [Ca^2+^]_i_ levels seen in TRPV6 transfected cells were possibly due to an effect of Fura-2 in buffering [Ca^2+^]_i_. Subsequent studies showed that TRPV6 is activated by a reduction in [Ca^2+^]_i_ concentration, and inactivated by higher [Ca^2+^]_i_ (Nilius et al., 2000; Yue et al., 2001). It should be mentioned that all previous studies were performed in cultured cells overexpressing TRPV6. Whether TRPV6 is constitutively open or is regulated in vivo was not clear. In this study, we generated a reporter fish line using the high-performance genetic calcium reporter GCaMP7a to measure intracellular Ca^2+^ activity in vivo. GCaMPs has been used for imaging intracellular Ca^2+^ activity in zebrafish neurons (Muto & Kawakami, 2013). This approach has alleviated the concern associated with Fura-2 and various cell culture systems overexpressing TRPV6. Our in vivo Ca^2+^ imaging results showed that the Trpv6 channel is constitutively open. This conclusion is supported by the facts that genetic deletion of *trpv6* reduced the [Ca^2+^]_i_ to undetectable levels and re-expression of Trpv6 restored the [Ca^2+^]_i_ levels in NaR cells. Addition of ionomycin did not result in further increase, indicating the endogenous [Ca^2+^]_i_ levels are high in these cells. More direct evidence came from the [Ca^2+^]_i_ dynamics analysis results. Within minutes after the addition of Trpv6 blockers GdCl_3_ or Ruthenium red, the levels of [Ca^2+^]_i_ were significantly reduced. When these drugs were washed out, the [Ca^2+^]_i_ signal levels returned to normal levels. It was thought that Ca^2+^ transporting epithelial cells, being continuously challenged by Ca^2+^ traffic from the apical side, maintain low levels of [Ca^2+^]_i_ using the cytosolic Ca^2+^ binding protein (Hoenderop et al., 2005). The in vivo imaging findings made in this study challenge this conventional view. We postulate that maintaining high [Ca^2+^]_i_ levels in these Ca^2+^ transporting cells is likely beneficial to the organism because it keeps these cells in differentiated state and functioning as Ca^2+^ transporting units. This idea is supported by the fact that treatment of zebrafish with the intracellular Ca^2+^ chelator BAPTA-AM promoted quiescent epithelial cells to proliferate.

The results in the present study suggest that Trpv6 mediates constitutive Ca^2+^ influx in epithelial cells to maintain their quiescent state and by continuously down-regulating IGF1 receptor-mediated Akt-Tor signaling. This is supported by the findings that 1) genetic deletion of Trpv6 or pharmacological inhibition of Trpv6-mediated Ca^2+^ uptake increased Akt and Tor signaling activity in an IGF1 receptor-dependent manner; and 2) re-expression of Trpv6 in the mutant cells suppressed Akt signaling. Our chemical biology screens and genetic studies identified PP2A as a key effector downstream of TRPV6/Trvp6. Inhibition of PP2A by two distinct inhibitors led to elevated Akt signaling and increased epithelial cell proliferation. Importantly, an IGF1 receptor inhibitor abolished these changes. Likewise, CRISPR/Cas9- mediated transient knockdown of PP2A catalytic subunits increased epithelial cell proliferation and Akt signaling. These in vivo findings, together with numerous biochemical studies showing that PP2A can dephosphorylate Akt (Perrotti & Neviani, 2013; Seshacharyulu et al., 2013), indicate that a [Ca^2+^]_i_-regulated PP2A isoform(s) likely acts downstream of TRPV6/Trpv6 and regulates the quiescent state in epithelial cells. This is in good agreement with a recent study in Drosophila showing that compromising PP2A activity delays cell cycle exit (Sun & Buttitta, 2015). The TRPV6-[Ca^2+^]_i_-PP2A-Akt signaling axis appears to be conserved in human colon cells because inhibition of TRPV6, [Ca^2+^]_i_ and PP2A all increased LoVo cell proliferation. The B regulatory subunits control substrate specificity and intracellular distribution of the PP2A holoenzymes and are encoded by a large set of genes classified into four families, i.e., B, B′, B″, and B″′ (Virshup & Shenolikar, 2009). Recent studies suggest that PR72/130 and PR70, 2 members of the B” family, possess 2 conserved EF hand motifs (termed EF1 and EF2) and increasing [Ca^2+^]_i_ increased the holoenzyme assembly and phosphatase activity (Kurimchak et al., 2013; Magenta et al., 2008). Another mechanism involves the action of [Ca^2+^]_i_-dependent m-calpain: m-calpain degrades PR72 and PR130 into a 45 kDa fragment (Janssens, Derua, Zwaenepoel, Waelkens, & Goris, 2009). This fragment, termed as PR45, is resistant to further degradation and exhibits enhanced PP2A activity. These two mechanisms may be related because the calpain-mediated proteolytic activation of PP2A depends on the EF hand integrity (Janssens et al., 2009). Future studies are needed to determine whether these mechanisms mediate TRPV6/Trpv6 action in epithelial cells.

Our findings linking the Trpv6-meiated Ca^2+^ uptake to the cellular quiescence regulation have important biomedical implications. Approximately 90% of human cancers arise in epithelial tissues and over-proliferation is one of the cancer hallmarks (Hanahan & Weinberg, 2011). The IGF-PI3K-AKT-mTOR pathway is one of the most frequently mutated signaling pathways in epithelial tissue-derived cancers, including colon and prostate cancers (Massoner, Ladurner-Rennau, Eder, & Klocker, 2010). The *IGF2* gene and *IRS2* gene are frequently gained in colon cancer (Cancer Genome Atlas, 2012) and have been proposed as a colorectal cancer “driver” oncogenes (Day et al., 2013). TRPV6 gene is frequently up regulated in prostate, colon, and other cancer tissues (V. Lehen’kyi, Raphael, & Prevarskaya, 2012; Prevarskaya, Skryma, & Shuba, 2018). At present, it is unclear whether the elevated TRPV6 expression promotes tumor growth or it is an adaptive response (V. Lehen’kyi et al., 2012; Prevarskaya et al., 2018). Knockdown or overexpression of TRPV6 in cultured cancer cells showed mixed effects in increasing/decreasing proliferation and/or apoptosis (Chow, Norng, Zhang, & Chai, 2007; V Lehen’Kyi, Flourakis, Skryma, & Prevarskaya, 2007; V. Lehen’kyi et al., 2012; Raphael et al., 2014; Skrzypski et al., 2016). Future studies are needed to clarify the role of TRPV6-[Ca^2+^]_i_-PP2A in prostate and colon cancer initiation and progression and its relationship to IGF1 receptor-PI3 kinase-AKT-mTOR signaling.

## Materials and methods

### Chemicals and reagents

All chemical reagents were purchased from Fisher Scientific (Pittsburgh, PA, USA) unless stated otherwise. The Phospho-Akt and phspho-S6 antibodies were purchased from Cell Signaling Technology (Beverly, MA, USA). Restriction enzymes were bought from New England Bio Labs (Beverly, MA, USA). PCR primers were synthesized by Life Technologies (Carlsbad, CA, USA). Cas9 protein was purchased from ToolGen (Seoul, South Korea). Rutheniuem red, and BAPTA-AM were purchased from MilliporeSigma (Burlington, MA, USA). Okadaic acid and Calyculin were purchased from Santa Cruz (Dallas, TX, USA) and Alomone Labs (Israel). BMS-754807 was purchased from Active Biochemicals Co (Hongkong, China).

### Zebrafish husbandry

Fish were raised following standard zebrafish husbandry guideline (Westerfield, 2000). Embryos were obtained by natural cross and staged following Kimmel at al. (Kimmel, Ballard, Kimmel, Ullmann, & Schilling, 1995). E3 embryo rearing solution (containing 0.33 mM [Ca^2+^]) was prepared as reported (Westerfield, 2000). Two additional embryo rearing solutions containing 0.2 mM [Ca^2+^] (i.e., normal [Ca^2+^] solution) or 0.001 mM [Ca^2+^] (i.e., low [Ca^2+^] solution) were made following previously reported formula (Dai et al., 2014). To inhibit pigmentation, 0.003% (w/v) N-phenylthiourea (PTU) was added in some experiments. All experiments were conducted in accordance with the guidelines approved by the University of Michigan Institutional Committee on the Use and Care of Animals.

### Generation of *trpv6*^*-/-*^ fish lines using CRISPR/Cas9

The sgRNA targeting *trpv6* (5’-GGGCTCGTTGATGAGCTCCG-3’) was designed using CHOPCHOP (http://chopchop.cbu.uib.no/). The sgRNA (30 ng/µl) was mixed with capped with Cas9 protein (700 ng/µl) and co-injected into *Tg(igfbp5a:GFP)* or wild-type embryos at the 1-cell stage as described (Xin & Duan, 2018). After confirming indels by PCR followed by hetero-duplex assay using a subset of F0 embryos, the remaining F0 embryos were raised to adulthood and crossed with *Tg(igfbp5a:GFP)* or wild-type fish. F1 fish were raised to the adulthood and genotyped. After confirming indels by DNA sequencing, the heterozygous F1 fish were intercrossed to generate F2 fish.

### Transient knockdown of *ppp2cs*

Three sgRNAs targeting *ppp2cs* were designed using CHOPCHOP (http://chopchop.cbu.uib.no/). Their sequences are: *ppp2ca*-sgRNA: 5’-GTTCCATAAGATCGTGAAAC-3’; *ppp2cb*-sgRNA: 5’-GAGCGTTCTCACTTGGTTCT-3’; *ppp2ca2*-sgRNA: 5’-GACGAAGGAGTCGAATGTGC-3’. sgRNAs (30 ng/µl) were mixed with Cas9 protein (700 ng/µl) and co-injected into *Tg(igfbp5a:GFP)* or wild type embryos at the 1-cell stage as reported (Xin & Duan, 2018). A subset of injected embryos were pooled, DNA isolated, and analyzed by PCR followed by hetero-duplex assays as reported (Chengdong Liu et al., 2018). After confirming the indels, the remaining injected embryos were used for experiments.

### Genotyping

To isolate genomic DNA, pooled embryos or individual adult caudal fin were incubated in 50 µl NaOH (50 mM) at 95 °C for 10 min and neutralized by adding 5 µl 1 M Tris-HCl (pH 8.0). PCR was performed using the following primers: *trpv6*-gt-f, 5’-TGACATTGTGTGTGTTTGTTGC-3’; *trpv6*-gt-r, 5’-GTGAAGGGCTGTTAAACCTGTC-3’; *trpv6*-HMA-f, 5’-GCAGCGGTGGCTTTAATGAAT-3’; *trpv6*-HMA-r, 5’-AAACCTGTCAATCAGAGCACAC-3’; *ppp2ca*-gt-f, 5’-TCACCATCAGTGCATGTCAATA-3’; *ppp2ca*-gt-r, 5’-CTCGATCCACATAGTCTCCCAT-3’; *ppp2cb*-gt-f, 5’-TGGATGATAAAGCGTTTACGAA-3’; *ppp2cb*-gt-r, 5’-ACGTTACACATTGCTTTCATGC-3’; *ppp2ca2*-gt-f, 5’-CTGATGGTTGTGATGCTGTTTT-3’; *ppp2ca2*-gt-r, 5’-CGGTTTCCACAGAGTAATAGCC-3’.

### Morphology analysis

Body length, defined as the curvilinear distance from the head to the end of caudal tail, was measured. Alizarin red staining was performed following a published protocol (Du, Frenkel, Kindschi, & Zohar, 2001). Images were captured with a stereomicroscope (Leica MZ16F, Leica, Wetzlar, Germany) equipped with a QImaging QICAM camera (QImaging, Surrey, BC, Canada).

### Whole-mount *in situ* hybridization, and immunostaining

For whole mount immunostaining or in situ hybridization analysis, zebrafish larvae were fixed in 4% paraformaldehyde, permeabilized in methanol, and analyzed as described previously (Dai et al., 2014). For double color *in situ* hybridization and immunostaining, mCherry mRNA signal was detected using anti-DIG-POD antibody (Roche), followed by Alexa 488 Tyramide Signal Amplification (Invitrogen). After in situ hybridization, the stained larvae were washed in 1X PBST and incubated with phosphorylated Akt antibody overnight at 4°C and then stained with a Cy3-conjugated goat anti-rabbit immunoglobulin G antibody (Jackson ImmunoResearch). Fluorescent images were acquired using a Nikon Eclipse E600 Fluorescence Microscope with PMCapture Pro 6 software.

### Plasmid and BAC constructs

The ORF of zebrafish Trpv6 were amplified by PCR and cloned into pEGFPN1 using the following primers: BglII-zftrpv6-F, 5’-atatAGATCTcgccaccATGCCACCCGCCATATC-3’; no stop-zftrpv6-ca-SalI-R, 5’-TACCGTCGACcaGAGAAACTTGAAATTggggcaatc-3’; Trpv6^D539A^ was engineered by site-directed mutagenesis using the following primers: zTrpv6_D539A_f, 5’-GGTCAGATTGCCTTGCCAGTGGA-3’; zTrpv6_D539A_r, 5’-TCCACTGGCAAGGCAATCTGACC-3’. Human TRPV6 ORF was sub-cloned into pEGFPN1 using the following primers: 5’-atatCTCGAGcgccaccATGGGTTTGTCACTG-3’; 5’-TACCGTCGACcaGATCTGATATTCC-3’. EGFP sequence in those vectors was replaced by mCherry sequence from pmCherry-C1 vector using the following primers: AgeI-mCherry-F, 5’-caACCGGTCGCCACCATGGTGAGCAAGGGC-3’; mCherry-NotI-stop-r, 5’-TCGCGGCCGCCTACTTGTACAGCTCGTCC-3’. Wild-type zebrafish Trpv6 and Trpv6D539A tagged with mCherry were then inserted into the *igfbp5aBAC* construct to replace the *igfbp5a* coding sequence from the start codon to the end of the first exon through homologous recombination as reported (C. Liu et al., 2017). The primers are: *igfbp5a-zTrpv56*-f, 5’-GTTTTGCCATTTCAAAGCTGGTGAAATAGGTGTTCTACAGTAGGACGATGCCACCCGCCATATCTGGTGAA-3’ and igfbp5a_frt-kan-rev, 5′-GTTTACTTTTGTCCCATATAAAACAAATACTACAAGTCAATAAAACATACCCGCGTGTAGGCTGGAGCTGCTTC-3′. The resulted BAC DNA was validated by sequencing. The validated BAC DNA and Tol2 mRNA were mixed and injected into 1-cell stage *trpv6*^*-/-*^; *Tg(igfbp5a:GFP)* embryos. The embryos were raised and analyzed. Cells co-expressing mCherry and GFP were identified and scored using a reported scoring system (Chengdong Liu et al., 2018).

### Generation of the *Tg(igfbp5a:GCaMP7a)* fish line

*GCaMP7a* DNA was cloned into pEGFPN1 to replace the EGFP sequence using the following primers: BamHI-HA-F, 5’-cgcggatccATGGCATACCCCTACGACG-3’; GCaMP7a-stop-NotI-R, 5’-atttgcggccgcTTACTTAGCGGTCATCATC-3’. *BAC* (*igfbp5a*:*GCaMP7a*) construct was generated following a published protocol (C. Liu et al., 2017). The following primers were used to amplify the GCaMP7a cassette sequence: *igfbp5a*_GCaMP7a_fw, 5′-GTTTTGCCATTTCAAAGCTGGTGAAATAGGTGTTCTACAGTAGGACGATGGCATACCCCTACGACGTGCCCGAC-3′ and igfbp5a_frt-kan-rev. The resulted *BAC (igfbp5a:*GCaMP7a) was validated by PCR using the following primers: igfbp5a-5UTR-fw, 5’-GAACGCTGTTCGCTTGAT-3’ and GCaMP7a-stop-NotI-R, 5’-ATTTGCGGCCGCTTACTTAGCGGTCATCATC-3’. The *BAC (igfbp5a:GCaMP7a)* DNA and *Tol2* mRNA were mixed and injected into zebrafish embryos at 1-cell stage. The F0 embryos were screened at 72 hpf by checking GCaMP7a responses to high or low [Ca^2+^] solutions. GCaMP7a-positive F0 embryos were raised and crossed with wild-type fish to obtain F1 individuals. F2 fish were generated by crossing F1 fish.

### Live GCaMP7a imaging

Zebrafish larvae were anesthetized using normal [Ca^2+^] embryo solution supplemented with 0.168 mg/ml tricaine. They were mounted in 0.3% low-melting agarose gel and immersed in 1 ml normal [Ca^2+^] solution. A Leica TCS SP8 confocal microscope equipped with the HC PL APO 93X/1.30 GLYC was used for imaging and LAS X and Image J were used for image analysis.

### Cell culture, transfection, Fura-2 imaging, and electrophysiology recording

HEK293 cells were cultured in DMEM supplemented with 10% FBS, penicillin and streptomycin in a humidified-air atmosphere incubator containing 5% CO2. Cells were plated onto 24 mm cover glass coated with L-polylysine and transfected with 0.3 µg of plasmid DNA using Lipofectamine 2000. Twenty-four hours after the transfection, the cover glass was mounted on an imaging chamber and washed with calcium-free Krebs-Ringer HEPES (KRH) solution (118 mM NaCl, 4.8 mM KCl, 1 mM MgCl_2_, 5 mM D-glucose, and 10 mM HEPES, pH=7.4). Fura-2 loading was performed following a published method (Kovacs et al., 2012). Successfully transfected cells were chosen by mCherry expression. Their cytosolic Ca^2+^ levels were recorded by an EasyRatio Pro system (PTI) at two different wavelengths (340 nm and 380 nm). The Fura-2 ratio (F340 /F380) was used to determine changes in intracellular [Ca^2+^]i. At least 50 cells were measured in each slide. Patch clamp recordings were performed at room temperature (Weissgerber et al., 2012). The internal pipette solution contained (in mM): Aspartate-Cs 145, NaCl 8, MgCl_2_ 2, HEPES 10, EGTA 10, Mg-ATP 2, pH 7.2. Normal external solution contained (in mM): NaCl 135, KCl 6, MgCl_2_ 1.2, HEPES 10, Glucose 12, pH 7.4, supplemented with 10 mM CaCl_2_ or 30 mM BaCl_2_. For measuring Ca^2+^ currents, cells were perfused with normal external solution at first and then switched to solutions as indicated. DVF solutions contains (in mM): NaCl 150, EDTA 10, HEPES 10, pH 7.4. Solutions with different concentration of Ca^2+^ contains (in mM): NaCl 150, HEPES 10, Glucose 12, pH 7.4, supplemented with 0 to 10 mM CaCl_2_ as indicated.

### Flowcytometry analysis

Human LOVO conlan cancer cells, obtained from ATCC, were cultured in DMEM/F12 supplemented with 10% fetal bovine serum (FBS), penicillin and streptomycin in a humidified-air atmosphere incubator containing 5% CO2. Cells were washed 3 times with serum-free medium (SFM) and starved in SFM for 12h. Drugs were then added together with 2% FBS - containing medium. Forty-eight hours later, cell cycle analysis was done using Attune Acoustic Focusing cytometer (Applied Biosystems, Life Technologies) after propidium iodide staining as described before (Chengdong Liu et al., 2018).

### Statistical analysis

Values are shown as Mean ± standard error of the mean (SEM). Statistical significance among experimental groups was determined using one-way ANOVA followed by Tukey’s multiple comparison test or student t-test. Chi-square test was used to analyze the association between two categorical variables. Statistical significances were accepted at *P* < 0.05 or greater.

## Acknowledgements

This work was supported by NSF grant IOS-1557850 and IOS-1755262. The funders had no role in study design, data collection and analysis, decision to publish, or preparation of the manuscript.

---

**Fig. S1.**
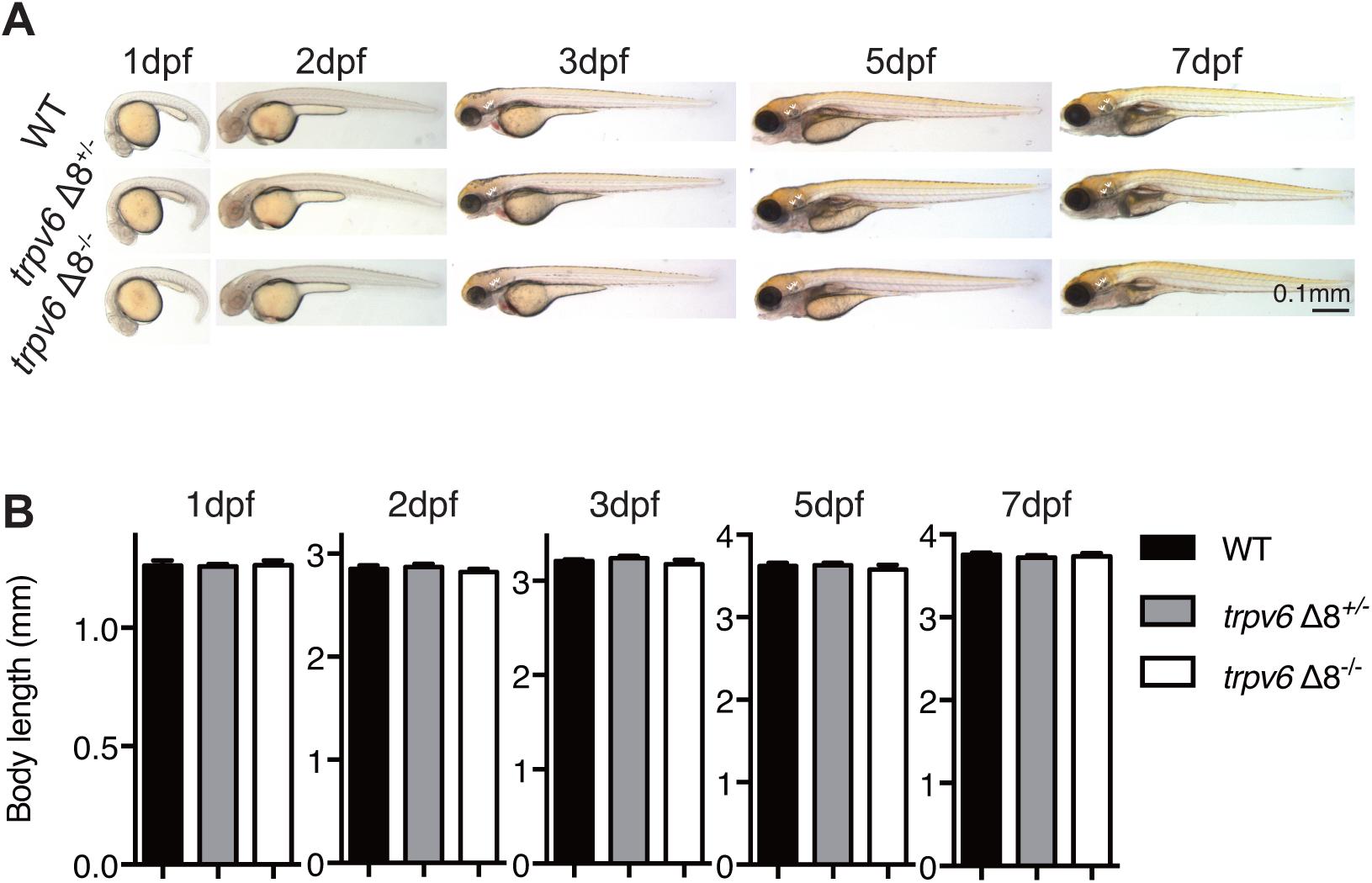
Morphology of *trpv6* mutant fish. (**A-B**) Gross morphology (**A**) and body size (**B**) of *trpv6Δ8*^*-/-*^; *Tg(igfbp5a:GFP)* fish and siblings. Values are Mean ± SEM, n = 7-23 fish. No statistical significance was found.

**Fig. S2.**
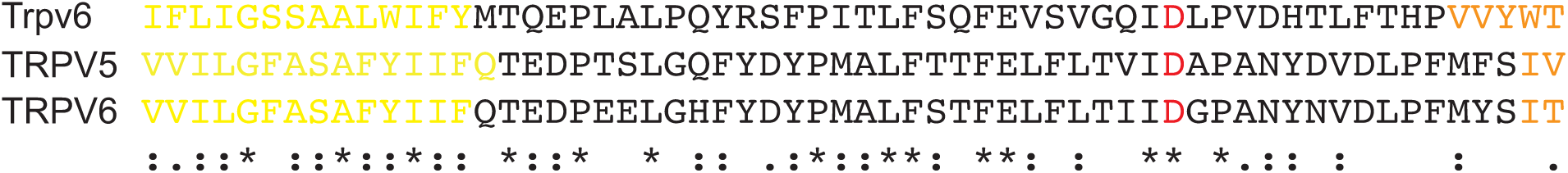
Sequence alignment of the zebrafish Trpv6, human TRPV5, and human TRPV6 pore region. Transmembrane domain 5 and 6 are indicated in yellow and orange letters, respectively. The ion pore region is indicated by black letters and the critical Asp residue (D) is labeled in red.

**Fig. S3.**
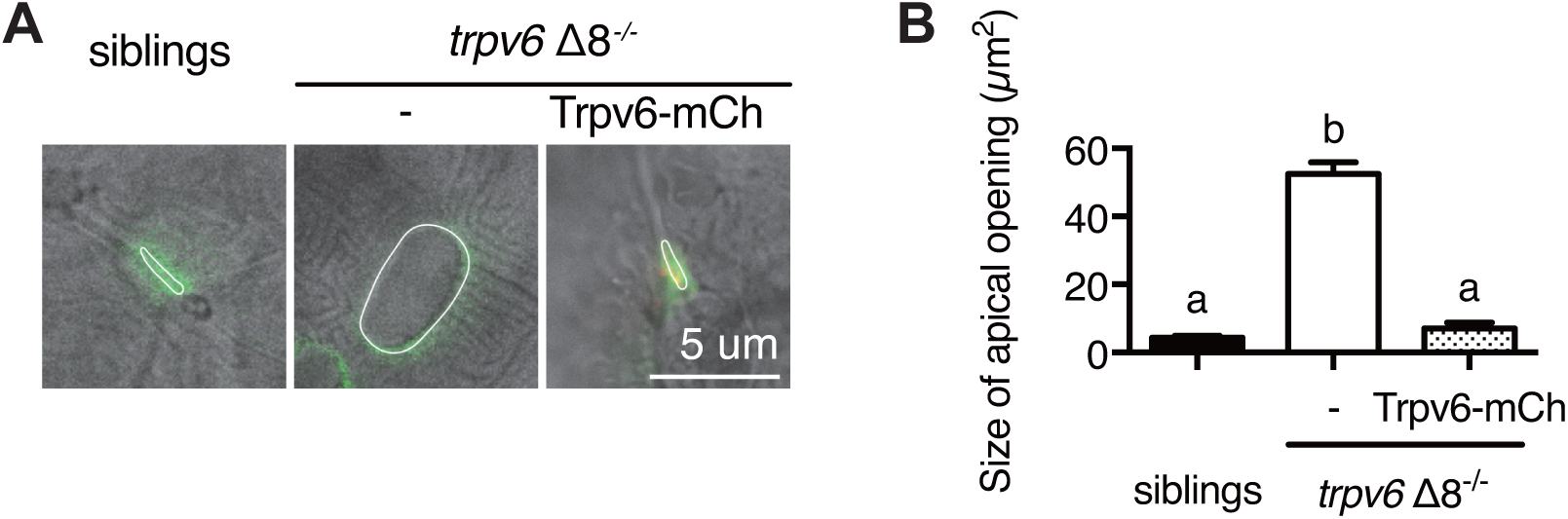
Genetic deletion of Trpv6 increases epithelial cell apical opening. (**A-B**) Progenies of a *trpv6Δ8*^*+/-*^*;Tg (igfbp5a:GFP)* intercross were injected with or without the *BAC (igfbp5a:Trpv6-mCherry)* DNA. At 3dpf, larvae were photographed followed by individual genotyping. Representative images are shown in (**A**). The apical opening of NaR cells were quantified and shown in (**B**). Data are Mean ± SEM, n = 8-42. Different letters indicate significant difference at *P* < 0.05, one-way ANOVA followed by Tukey’s multiple comparison test.

**Fig. S4.**
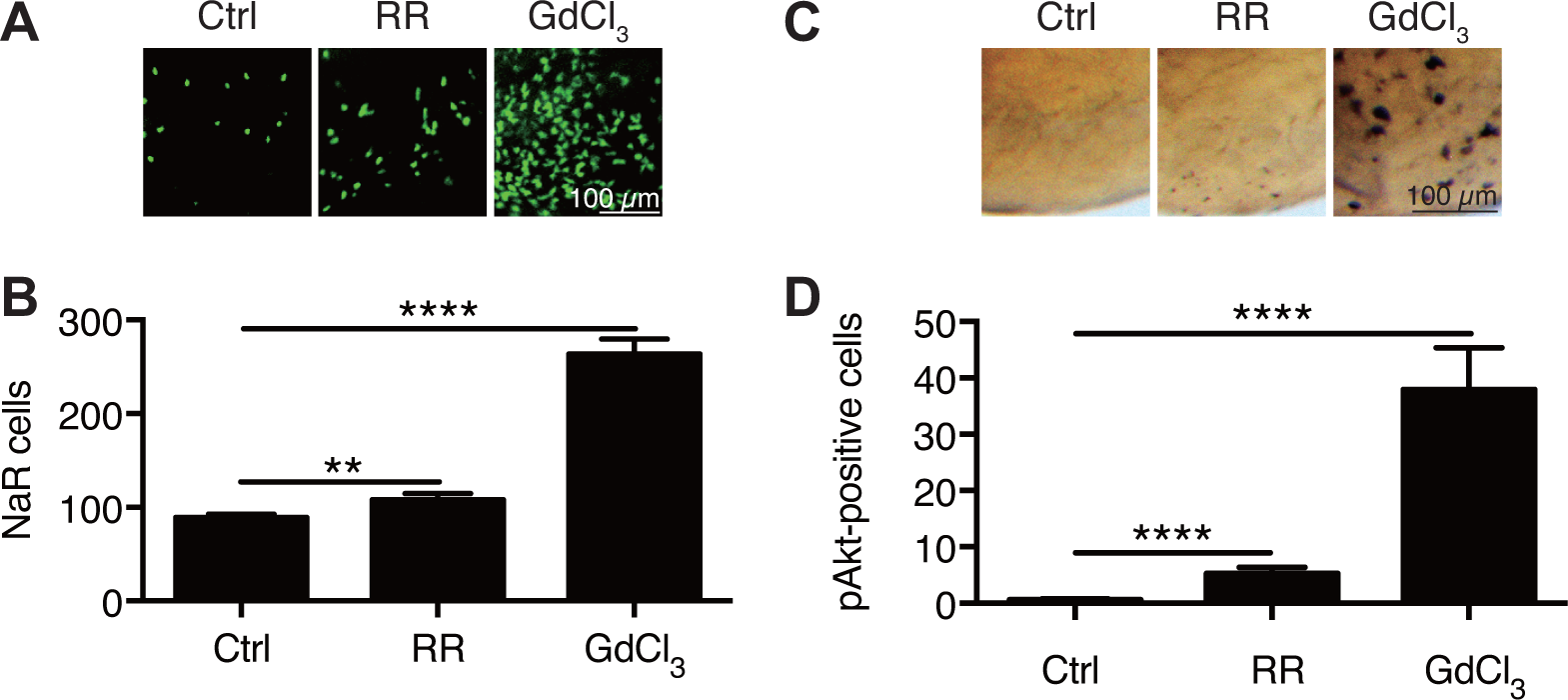
Inhibition of Trpv6 increases epithelial cell proliferation and Akt signaling. (**A-B**) *Tg(igfbp5a:GFP)* fish were treated with Ruthenium Red (2 µM) or GdCl_3_ (100 µM) from 3 to 5 dpf. Representative images are shown in (**A**). NaR cell numbers were quantified and shown in (**B**). (**C-D**) Wild type fish were treated with Ruthenium Red (30 µM) or GdCl_3_ (100 µM) from 3 to 4 dpf and analyzed by immunostaining using an anti-phospho-Akt antibody. Representative images are shown in (**C**) and quantified results shown in (**D**). **, **** indicate *P* < 0.01 and 0.0001 by unpaired t-test.

**Fig. S5.**
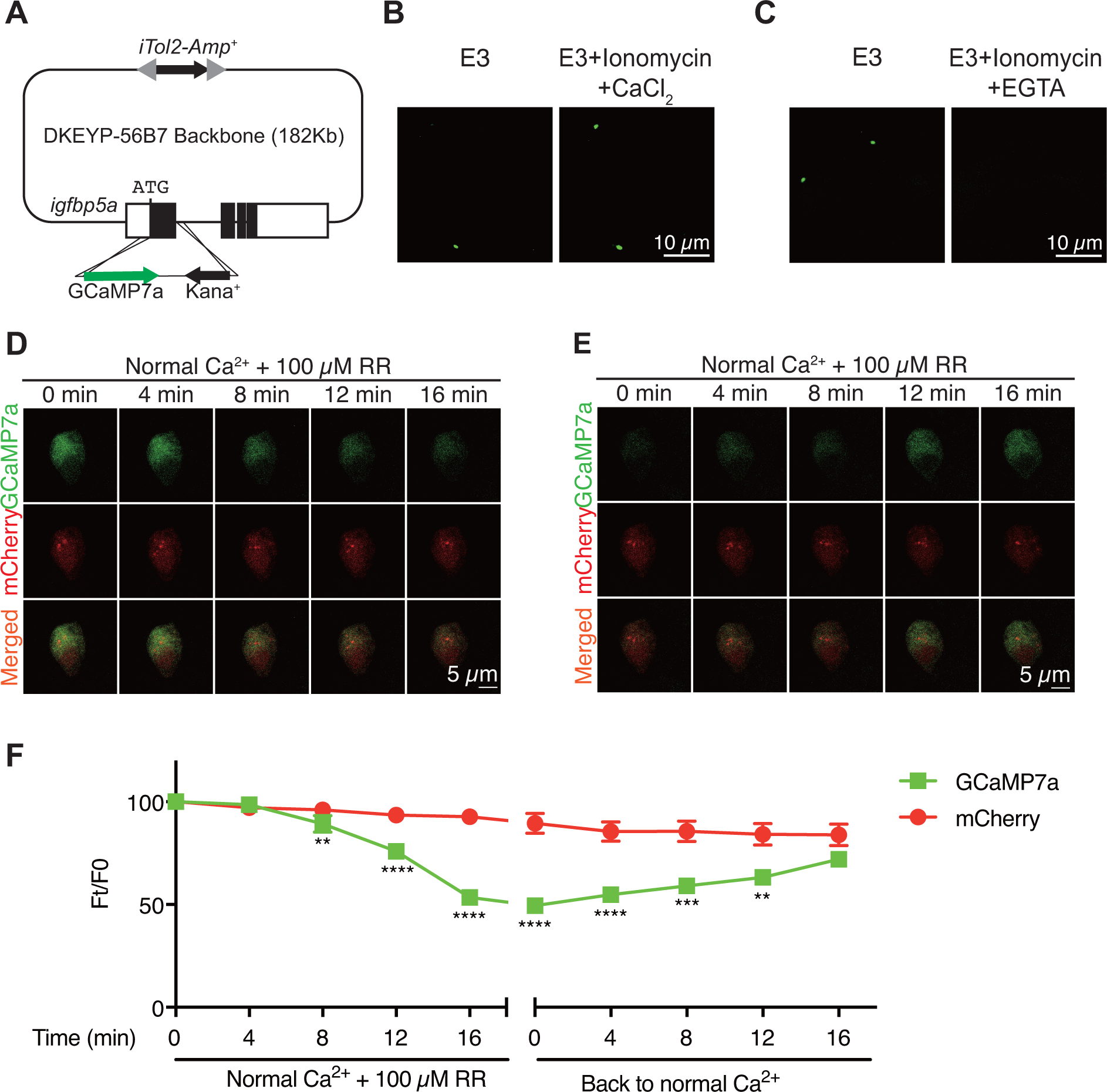
Generation and validation of *Tg(igfbp5a:GCaMP7a)* fish. (**A**) Schematic diagram showing the *BAC(igfbp5a:GCaMP7a)* construct engineered. Filled boxes indicate *igfbp5a* ORF and open boxes indicate its UTRs. The *iTol2* cassette and GCaMP7a reporter cassette were introduced into DKEYP-56B7 by homologous recombination. The *igfbp5a* sequence from the start codon to the end of first exon was replaced by the GCaMP7a cassette. (**B-C**) Embryos injected with *BAC(igfbp5a:GCaMP7a)* DNA were raised in E3 embryo solution and imaged at 3 dpf before and after the addition of the indicated chemicals (Ionomycin: 5 µM, CaCl_2_: 10 mM, EGTA: 10 mM). (**D-E**) Time-lapse images of 3 dpf larvae were taken after the addition (**D**) and removal (**E**) of 100 µM Ruthenium red (RR) at the indicated time points. Fluorescence change of GCaMP7a (green) and mCherry (red) were quantified and shown in (**F**). Mean ± SEM, n = 8. **,***, and **** indicate *P* < 0.01, *P* < 0.001, *P* < 0.0001 by multiple t-tests.

**Fig. S6.**
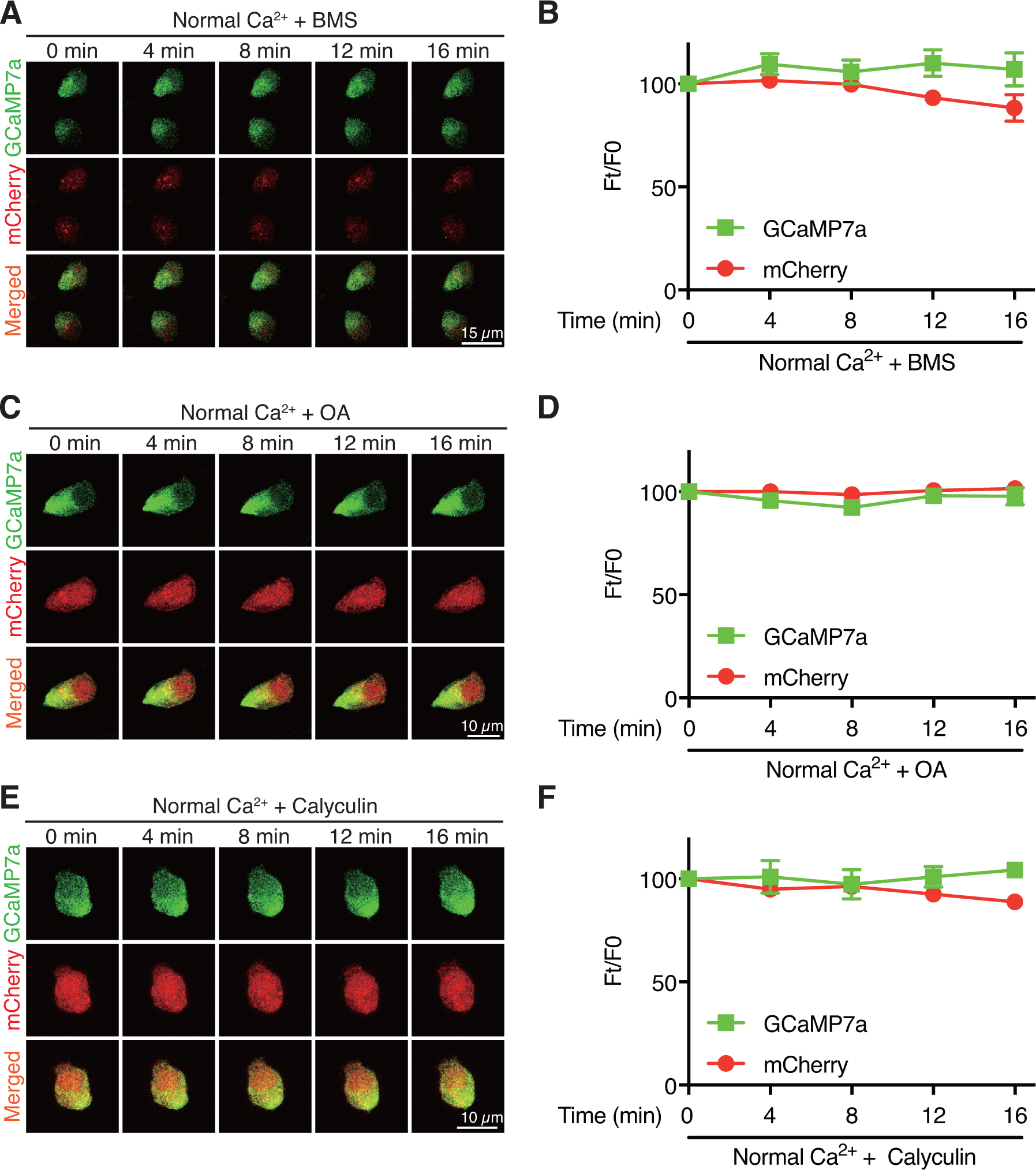
Inhibition of IGF1 receptor or PP2A does not change [Ca^2+^]_i_ in NaR cells. Time-lapse images of 3 dpf *Tg (igfbp5a:GCaMP7a)* larvae at the indicated time points after adding 0.3 µM BMS-754807 (**A, B**), 1 µM Okadaic acid (OA) (**C, D**) or 0.1 µM Calyculin (**E, F**). Changes in GCaMP7a (green) and mCherry (red) signal intensity were quantified. Representative images are shown in (A, C, and E) and quantified results are shown in (**B, D**, and **E**). Mean ± SEM, n = 3. No significance was found by two-way ANOVA followed by Dunnett’s multiple comparisons test.

**Fig. S7.**
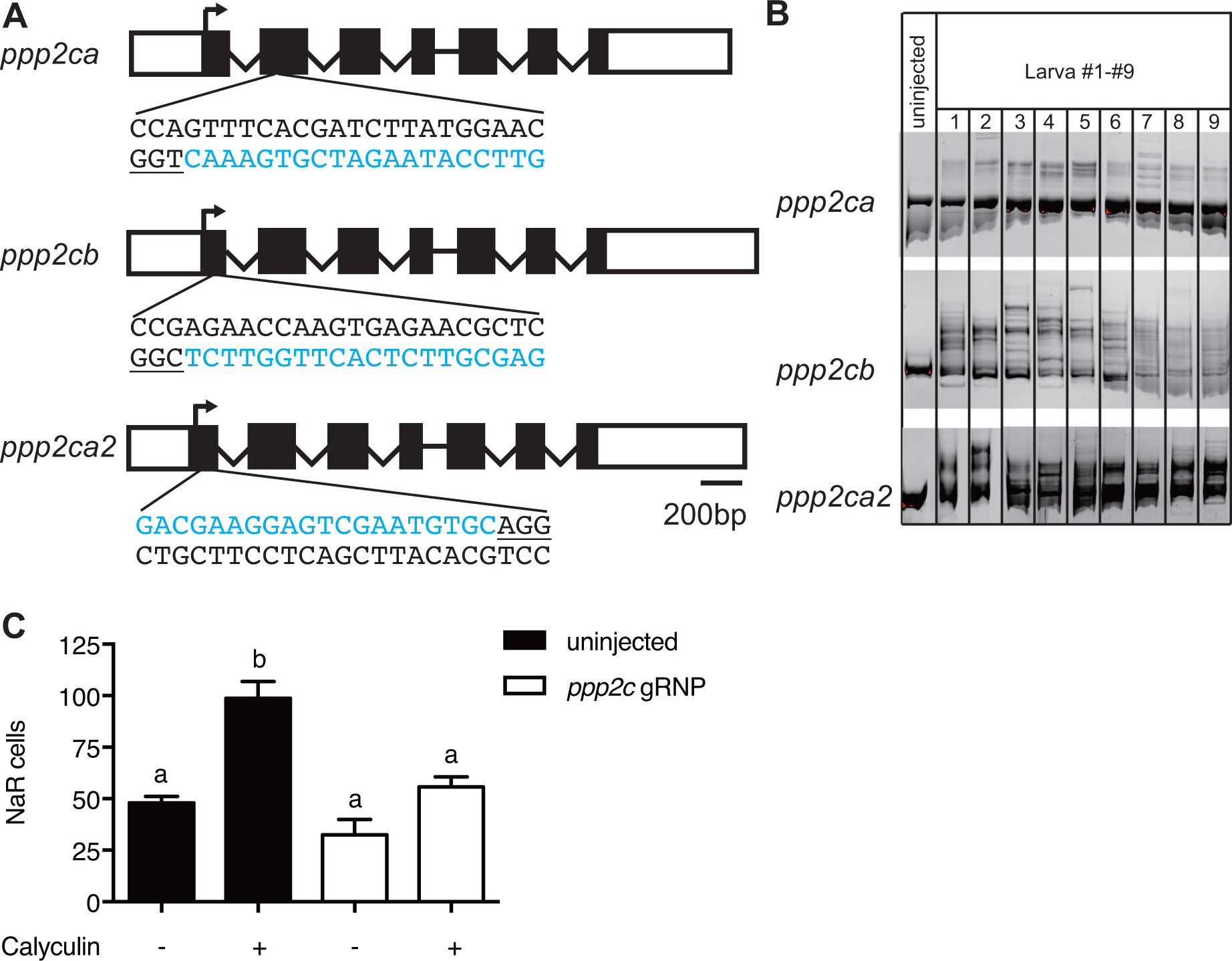
Transient knockdown of *pp2a* catalytic subunit genes. (**A**) Schematic diagram and guide RNA targeting sites. The target sites are labeled by blue letters and the PAM motif is underlined. (**B**) Embryos injected with gRNAs and Cas9 protein were raised to 1 dpf. Each of them was lysed and analyzed by PCR followed by hetero-duplex motility assay. (**C**) Embryos injected with gRNAs and Cas9 protein were raised to 3 dpf and treated with Calyculin or vehicle. Values are Mean ± SEM, n = 7-15. Different letters indicate significant differences between groups (*P* < 0.05, one-way ANOVA followed by Tukey’s multiple comparison test).

